# The Infant Brainstem - A Multimodal Multiscale Postmortem Imaging Pipeline

**DOI:** 10.64898/2026.08.28.744504

**Authors:** Caroline Magnain, Jean Augustinack, Chris Clickner, William Ammon, Nathan Ngo, Andre van der Kouwe, Brian L. Edlow, Robin Haynes, Luiz Fernando Ferraz da Silva, Hannah Kinney, Lilla Zöllei

## Abstract

Brainstem disorders in human infants - including sudden infant death syndrome (SIDS), the leading cause of postnatal infant mortality in the United States - are characterized by cellular and molecular abnormalities that conventional neuroimaging cannot detect. A substantial challenge arises from the fact that the immature myelination of the infant brain severely degrades MRI contrast, leaving the discrete nuclei and white matter tracts of the brainstem poorly resolved at the scale where pathology occurs. We introduce a postmortem imaging pipeline that bridges this gap by integrating four spatially registered modalities: whole-brain magnetic resonance imaging (MRI) (550 *µ*m), brainstem-specific MRI (150 *µ*m), polarization-sensitive optical coherence tomography (PSOCT, 10 *µ*m), and histology with immunohistochemistry (1.88 *µ*m). Our central finding is that PSOCT provides excellent tissue contrast independent of myelination state - directly overcoming the principal limitation of MRI in the infant brain - while enabling three-dimensional visualization of nuclei and tracts at resolutions 15 to 55 times finer than MRI alone. Histology provides cellular-level ground truth and validates the optical contrasts. Applied here to a normative 34-day-old infant brainstem, this pipeline establishes a generalizable framework for studying infant brainstem neuroanatomy in three dimensions, with particular relevance to disorders such as SIDS where gross anatomy is intact but cellular abnormalities remain the target of investigation.

## 1 Introduction

Brainstem disorders of the human infant represent a heterogeneous and clinically challenging group of pathologies spanning a wide range of severity. At one end are major structural malformations detectable by conventional neuroimaging, including pontocerebellar hypoplasia type 2 (Barth et al., 2007; Ekert et al., 2016), Dandy-Walker malformation (Alves et al., 2023), Arnold-Chiari malformation (Fons and Jnah, 2021), and brainstem disconnection syndrome (Duffield et al., 2009). At the other end — and more difficult to study — are disorders in which gross anatomy is entirely intact, but cellular, biochemical, or molecular abnormalities drive the pathology. These include Congenital Central Hypoventilation Syndrome (Weese-Mayer et al., 2021), RETT syndrome (Glaze, 2005), Prader-Willi syndrome (Yamada et al., 2022), and sudden infant death syndrome (SIDS) (Kinney and Haynes, 2019; Kinney et al., 2009), defined as the sudden and unexpected death of a seemingly healthy infant that remains unexplained after a thorough review of clinical history, anatomic and histologic autopsy, and death scene investigation (Goldstein et al., 2019). SIDS remains the leading cause of postnatal infant mortality in the United States today (Pretorius and Rew, 2019).

Understanding this second category of brainstem disorders requires imaging tools capable of resolving individual nuclei and white matter tracts — the small-scale structures where pathology is expressed. This is an intrinsically difficult problem. The brainstem spans between the cerebrum and spinal cord, comprising dozens of functionally distinct gray matter nuclei and adjacent white matter tracts tightly packed in a small anatomic space with complex three-dimensional organization. Detecting cellular abnormalities in this context demands both high spatial resolution and reliable tissue contrast.

Magnetic resonance imaging, introduced clinically in the early 1980s, has been central to brainstem research both in living individuals and the autopsy brain. Technological advances — including ultra-high field MRI (Sclocco et al., 2018) and extended *ex vivo* acquisition protocols (Augustinack et al., 2005; Edlow et al., 2019) — have substantially improved resolution and contrast. Yet even the finest current MRI approaches have not reached the 0.2–0.5 *µ*m resolution of the light microscope in 2D sections, the scale at which neuroanatomy and neuropathology are ultimately defined. More critically for infant imaging, MRI contrast depends substantially on myelin. Because white matter tracts in the infant brain are at varying and incomplete stages of myelination, they contain proportionally more water than in the fully myelinated adult brain, producing lower and often inverted contrast between gray and white matter. This is not a limitation that higher field strength or longer acquisition time can fully overcome — it is a fundamental property of the tissue. As a result, the discrete nuclei and tracts most relevant to brainstem pathology are precisely those that MRI resolves least reliably in the infant.

Optical imaging methods offer complementary strengths. Three-dimensional polarized-light imaging (PLI) (Axer et al., 2011; Larsen et al., 2007; Reckfort et al., 2015) exploits the birefringence of myelinated fibers to map fiber tract orientations of the whole brain at 100 *µ*m and higher axial resolution and down to a few microns in lateral resolution, and light-sheet fluorescence microscopy (LSFM) (Costantini et al., 2023; Stelzer et al., 2021; Ueda et al., 2020) coupled with tissue clearing, such as CLARITY (Chung et al., 2013) or SHORT (Pesce et al., 2022), can visualize fluorescently targeted features at micrometer resolution. However, both require the tissue to be sectioned prior to imaging, which introduces distortions and tears that make accurate three-dimensional reconstruction difficult. Optical coherence tomography (OCT) and polarization-sensitive optical coherence tomography (PSOCT) (Huang et al., 1991) avoid this limitation by imaging intact tissue blocks before sectioning, relying solely on the intrinsic optical properties of the tissue and the birefringence of the myelin sheath, respectively (Chang et al., 2022; Liu, Ammon, Siless, et al., 2021; Magnain et al., 2014; Wang et al., 2016, 2018). Critically, PSOCT contrast does not depend on the degree of myelination, making it well suited to the in-depth imaging of the infant brain.

Wang et al. (Wang et al., 2025) used PSOCT on slabs of cerebrum of developing brains, from 0 to 5 years old, showing the impact of myelination on the scattering coefficient, a contrast obtained by PSOCT imaging. The younger brains, less than 15 months of age, have a lower scattering coefficient in the white matter which the authors attributed to largely unmyelinated fibers (Deoni et al., 2011; Dubois et al., 2014; Hasegawa et al., 1992). Contrary to the previous methods, (PS)OCT images the tissue block prior to sectioning which greatly limits the distortions of the tissue and improves its 3D reconstruction and alignment to other modalities. This property is extremely useful for our 3D multimodal imaging effort.

To investigate the morphology of the developing infant brainstem at scales and resolutions previously inaccessible, we developed a multimodal postmortem imaging pipeline that spatially registers three complementary modalities of four spatial resolutions in a common three-dimensional coordinate system (Figure 1). Whole-brain and brainstem-specific MRI provide anatomical context and a bridge to *in vivo* measurements. PSOCT captures fine structural detail on intact tissue blocks across the full brainstem volume, independent of myelination state. Histology and immunohistochemistry — including stains for healthy neurons, myelinated fibers, and serotonergic neurons — provide cellular-level ground truth and a basis for validating the optical contrasts. Together, these modalities span four orders of resolution magnitude, from 550 *µ*m to 1.88 *µ*m (Table 1), unified in a shared spatial framework that enables multiscale analysis no single modality could support alone. We demonstrate this pipeline on a normative infant brainstem, establishing a proof-of-concept foundation for future application to cases where brainstem pathology — including SIDS — may be expressed at the cellular rather than the gross anatomical scale.

**Figure 1:**
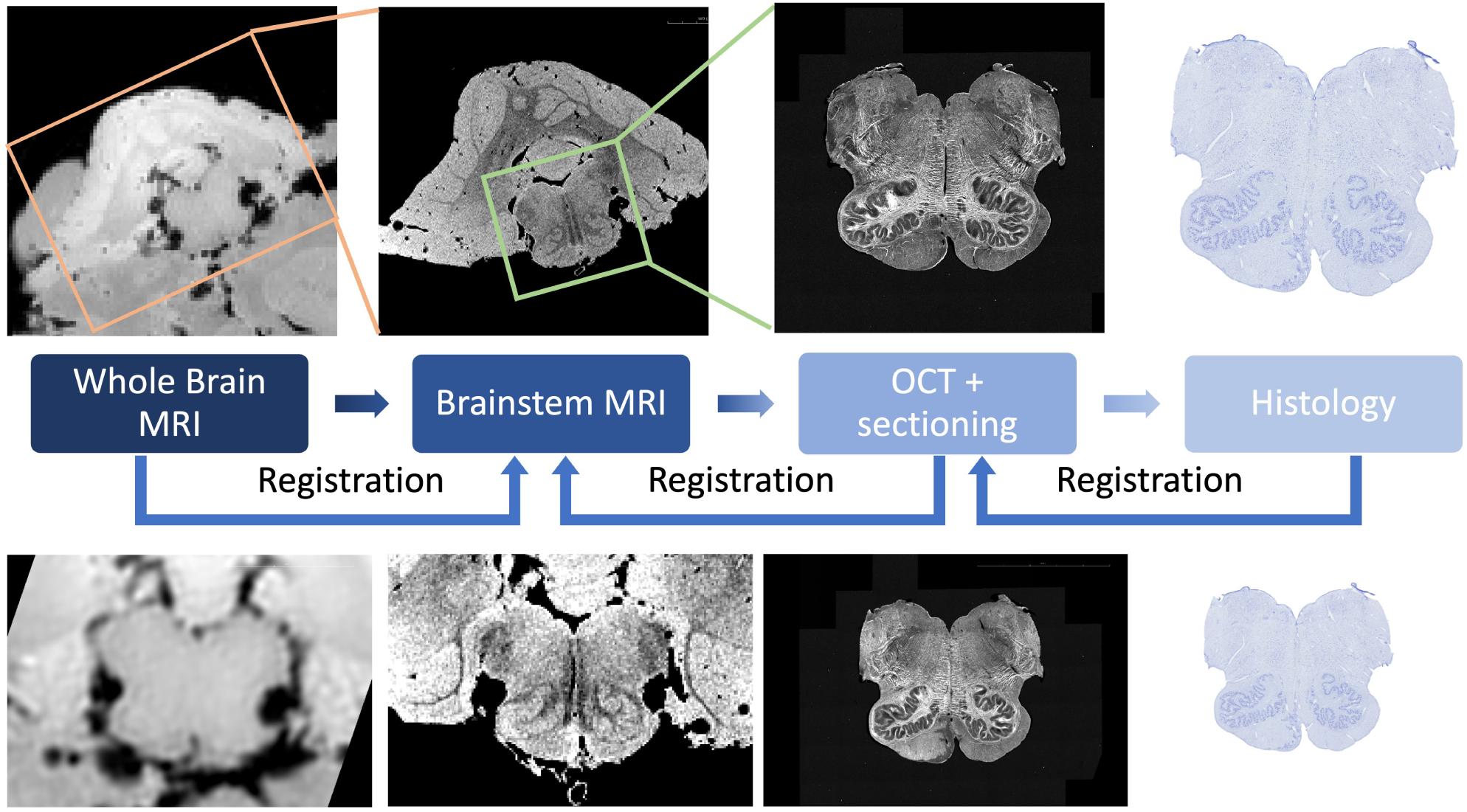
Overview of the postmortem infant brainstem imaging pipeline: whole brain MRI, brainstem MRI, OCT and histology. The orange box in the whole brain MRI shows the region blocked for the brainstem MRI, including the cerebellum and the brainstem. The green box shows the blocked region imaged by OCT, in this instance the medulla. The images in the bottom row zoom in on the same region of interest imaged at the 4 stages of the pipeline.

**Table 1:** Details of imaging modalities and their resolutions.

| Imaging Modality | Resolution | Details |
| --- | --- | --- |
| Whole brain MRI | 550 $\mu\text{m}$ | 3 T |
| Brainstem MRI | 150 $\mu\text{m}$ | 7 T |
| PSOCT | Projection volumes: 10 x 10 x 100 $\mu\text{m}^3$<br>$\mu_s$ volume: 10 and 20 $\mu\text{m}$ isotropic | AIP, MIP, retardance,<br>optic axis orientation, $\mu_s$ |
| IHC / Histology | 1.88 $\mu\text{m}$ (series evenly spaced every 200 $\mu\text{m}$) | Nissl, MBP, TPH2 |

AIP: Average Intensity projection, MIP: Maximum Intensity Projection, $\mu_s$ : scattering coefficient, IHC: immunohistochemistry, MBP: Myelin Basic Protein, TPH2: Tryptophan Hydroxylase 2.

## 2 Methods and Materials

### 2.1 Pipeline Overview

Our imaging pipeline integrates three modalities at four resolutions chosen to complement each other across scale and contrast mechanism (Table 1). Whole-brain MRI establishes anatomical context and provides a bridge to *in vivo* measurements. Brainstem-specific MRI increases resolution and contrast within the region of interest. PSOCT images intact tissue blocks at micrometer resolution using intrinsic optical properties of the tissue, independent of myelination state. Histology and immunohistochemistry provide cellular-level ground truth and a basis for interpreting and validating the optical contrasts. We spatially register all modalities into a common coordinate system, enabling multiscale information to be visualized and analyzed in a shared three-dimensional space. The pipeline is applied here to the brainstem of a normative 34-day-old infant as a proof-of-concept; the tissue showed no macroscopic or histological abnormalities, making it an appropriate reference case for pipeline validation.

### 2.2 Tissue Sample

The tissue imaged for this work was a medulla, pons, and midbrain from a full-term, 34-day-old male infant (cause of death identified as SIDS), sourced from the Department of Pathology, University of São Paulo, São Paulo, Brazil, where all consent procedures were performed in accordance with local study protocols and standard operating procedures. The clinicopathologic diagnosis was SIDS. While microscopic cellular and myelin defects have been reported in small subsets of SIDS cases (Becker, 1990; Kinney et al., 2025), a proportion of SIDS cases in each cohort present without microscopic deficits; this case fell into the latter category, making it suitable as a normative reference for pipeline development.

Tissue procurement was coordinated by local autopsy technicians and assessed by a neuropathologist (LFFS). The postmortem interval was 6.5 hours and brain removal followed an established protocol (Burton and Rutty, 2010). The specimen was fixed in neutral buffered 10% formalin for 129 days, with weekly formalin changes for the first month to ensure adequate fixation, then transferred to the Athinoula A. Martinos Center for Biomedical Imaging at Massachusetts General Hospital.

### 2.3 MRI Imaging

Prior to imaging, the brain sample was transferred to Fomblin (perfluoropolyether, Kurt J. Lesker Company, PA, USA) for at least one week to remove air bubbles trapped in the cortical folds. Imaging in Fomblin eliminates the background signal around the sample and reduces magnetic susceptibility artifacts. The sample was secured inside the coil with padding to minimize motion artifacts induced by vibrations during gradient switching.

Two MRI datasets were acquired. For whole-brain imaging, we used a 3 T Siemens TIM Trio with a 32-channel head coil, acquiring 550 *µ*m isotropic voxel resolution scans using a multi-echo FLASH (MEF) sequence (TR = 40 ms; flip angles *α* = 10°, 20°, 30°; TE = 4.96, 12.28 ms; acquisition time 35:31 min:s per flip angle). We share the volumes on the DANDI platform (https://about.dandiarchive.org/) and visualize them using Neuroglancer (Maitin-Shepard and Neuroglancer contributors, 2025): Whole Brain MRI. The brainstem and cerebellum were then dissected and imaged at 7 T with a solenoid coil (ID 68 mm) at 150 *µ*m isotropic resolution using MEF (TR = 25 ms; *α* = 3°, 5°, 10°, 15°, 20°, 30°, 40°; TE = 6.76, 13.18 ms; acquisition time 1:23:38 h:min:s per flip angle). We share the volumes on the DANDI platform and visualize them using Neuroglancer: Brainstem MRI. Parameter maps — including proton density and T1 relaxation times — were generated from the multi-echo, multi-flip angle data using FreeSurfer (Deoni et al., 2005; Fischl et al., 2004). We share the data on the DANDI platform and visualize them using Neuroglancer: parameter maps for the whole brain and brainstem. Diffusion MRI was also acquired for both the whole brain and brainstem, but is not discussed here in order to constrain the scope of this paper.

### 2.4 Optical Coherence Tomography

We selected PSOCT as the intermediate-scale modality for two reasons. First, unlike PLI and LSFM, PSOCT images intact tissue blocks before sectioning, greatly limiting tissue distortion and facilitating three-dimensional reconstruction and alignment to other modalities. Second, PSOCT contrast derives from the intrinsic optical properties of the tissue — specifically the birefringence of the myelin sheath — rather than from the degree of myelination *per se*. This makes it effective in the infant brain at stages when myelin content is low and variable, precisely the condition where MRI contrast is most compromised.

#### 2.4.1 Tissue Preparation and Acquisition

To fit within the imaging system, we cut the brainstem into three blocks (Figure 2A): the caudal medulla (block 1), the rostral medulla and caudal pons (block 2), and the rostral pons and midbrain (block 3). Cuts were placed where brainstem features vary slowly — in the middle of the medulla and middle of the pons — to preserve the most rapidly changing transitional regions between structures intact. Meninges and pia were removed to facilitate clean vibratome sectioning. Each block was embedded in melted oxidized agarose. Block 3, unlike the other two, was imaged across two sessions due to its size; one session covered the rostral pons and a little bit of the mibrain, and the other the rest of the midbrain region.

**Figure 2:**
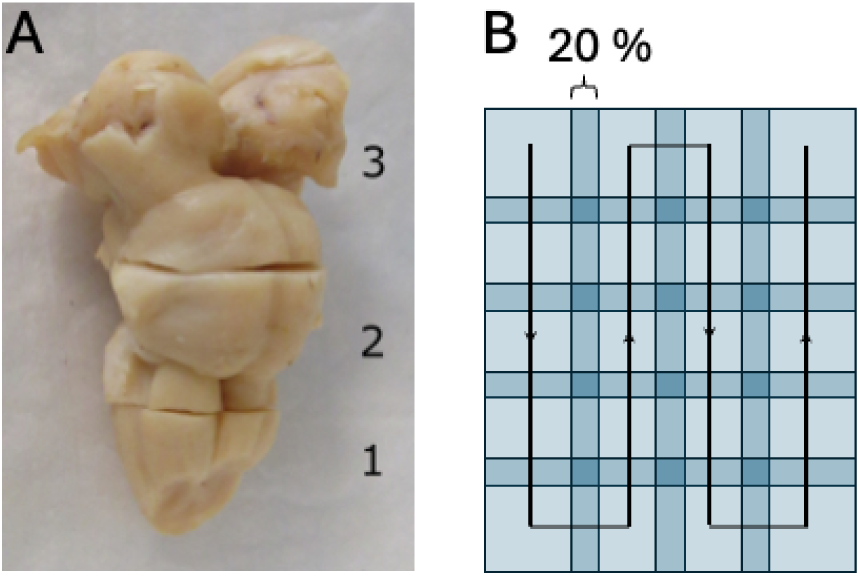
A) Brainstem blocking for OCT imaging. 1) Caudal medulla, 2) rostral medulla and caudal pons, 3) rostral pons and midbrain. B) Serpentine scheme for the OCT acquisition showing the 20% overlap.

We used a commercial spectral-domain PSOCT system centered at 1300 nm (TEL220PS, Thorlabs, Inc.) (Liu, Ammon, Jones, et al., 2021; Wang et al., 2025), yielding an axial resolution of 3.5 *µ*m in tissue and an imaging depth of 2.6 mm. A scan lens (OCT-LSM03, Thorlabs, Inc.) provided 10 *µ*m lateral resolution over a 3.5 × 3.5 mm^2^ field of view and a confocal parameter of 300 *µ*m. XYZ motorized stages translated the tissue sample; a vibratome performed sectioning. The acquisition sequence for each block was as follows:

1. Section the sample with the vibratome to ensure a flat surface
2. Acquire OCT images across the entire blockface in a serpentine-like manner using the XY stages, with a 20% overlap between adjacent tiles (Figure 2B)
3. Section two 50-*µ*m-thick slices and collect them in sequential order for histology or immunohistochemistry
4. Repeat steps 2 and 3 until the whole tissue block has been imaged.

The entire procedure was automated and controlled by our custom laboratory software.

#### 2.4.2 OCT Processing

##### Tile processing

Each tile was processed individually to produce four contrast volumes. The average intensity projection (AIP) was computed as a proxy for tissue attenuation, primarily due to scattering mixed with surface backscattering in the brain; this shows contrast between brainstem nuclei and tracts. The maximum intensity projection (MIP) emphasizes features with high refractive index contrast, including myelin and neurons; fiber tracts running perpendicular to the incident light appear bright, while those running parallel appear dark. Retardance — the cumulative phase shift between orthogonal polarization components — reflects myelin density, with brightness proportional to density. The optic axis orientation encodes the in-plane fiber direction, weighted by retardance to highlight myelinated regions; color represents orientation as shown in Figure 3. These four contrasts are projections over a depth of 300 *µ*m.

**Figure 3:**
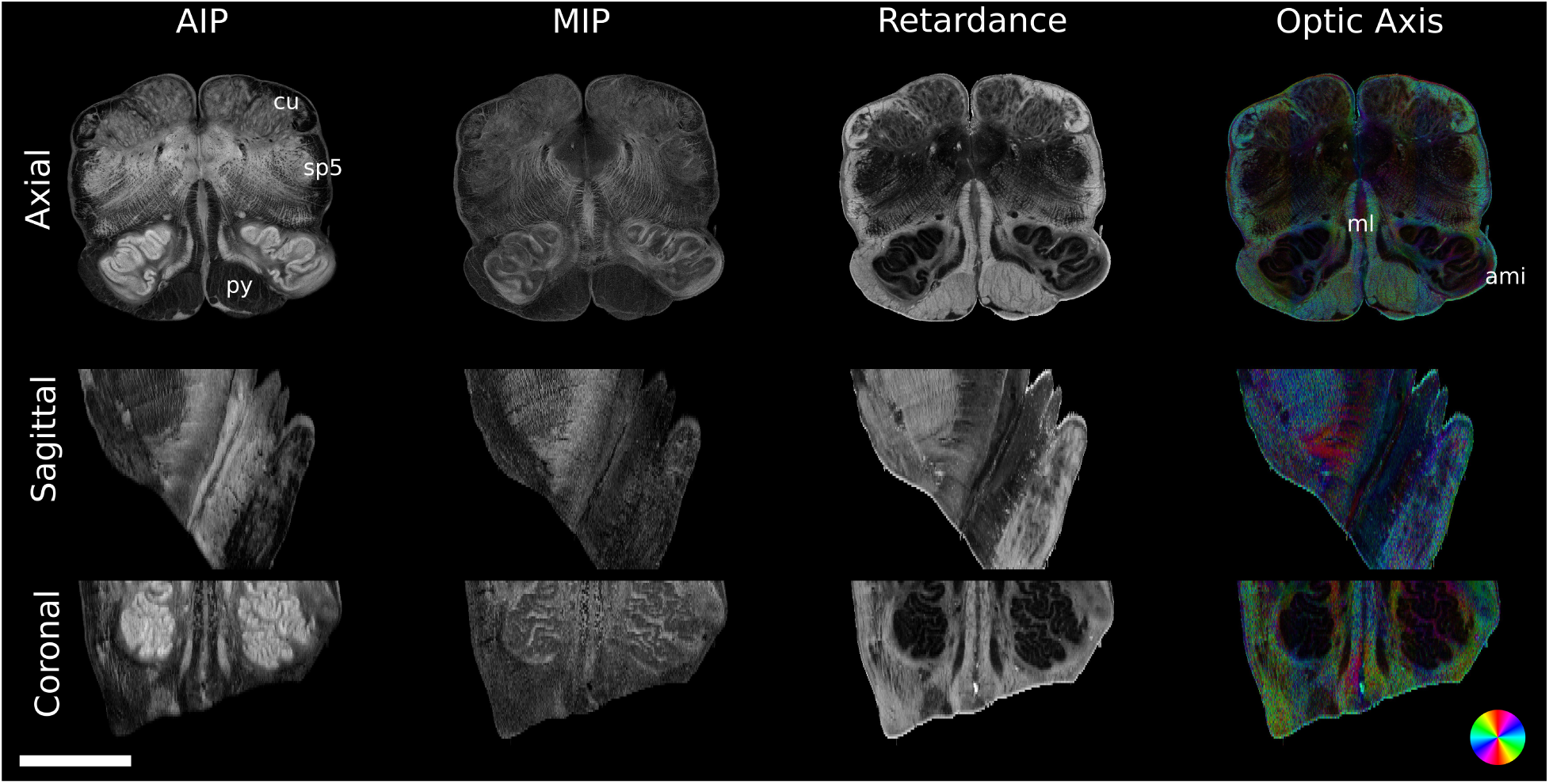
OCT projection volume acquired on the caudal medulla block. The figure shows the AIP, MIP, retardance and optic axis from the three standard orthogonal views: axial, sagittal and coronal. The scale bar is 5 mm and the color wheel shows the in-plane fiber orientation represented by the optic axis volume. ami: amiculum, cu: cuneate tract, ml: medial lemniscus, py: pyramid, sp5: spinal trigeminal tract.

The depth-dependent scattering coefficient *µ*_s_ was then computed for each tile following the method of Vermeer et al. (Vermeer et al., 2014) after being corrected for systematic geometric distortions due to our optical system and the roll-off of the detection camera:

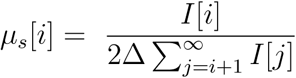

where *i* is the pixel index in depth and Δ is the axial pixel size. In our study we sum the intensity to the last saved voxel in depth, down to at least 600 *µ*m below the surface at least, reaching the noise level for a fixed tissue.

##### Stitching

Tiles were spatially aligned using the AIP contrast and the 20% inter-tile overlap, and slices were reconstructed independently using Fiji (Preibisch et al., 2009). Median XY coordinates across all slices were used to reconstruct projection volumes for all four contrasts.

Moreover, in order to reconstruct an isotropic volume, to correct for small inaccuracies in Z-stage positioning and vibratome sectioning unevenness, tiles were aligned in the Z direction by maximizing the 2D cross-correlation between various depths of adjacent tiles at each XY location, using a custom Matlab script. Only tiles whose surface represents at least 50% of the tissue are accounted for in this computation.

We created the individual optical volumetric slices. Each of these 3D slices were reconstructed in intensity (dBI) in their native voxel size, 10 x 10 x 2.5 *µ*m ^3^ using the XY coordinates determined previously. The scattering coefficient volumes were computed at isotropic pixel sizes of 10 *µ*m and 20 *µ*m. Gray matter, containing cells, presents lower *µ*_s_; white matter tracts and cranial nerves present higher *µ*_s_, with myelinated fibers showing the highest values. Finally, the full scattering coefficient isotropic volumes were reconstructed.

### 2.5 Histology and Immunohistochemistry

Histology and immunohistochemistry serves two roles in this pipeline: it provides cellular-level ground truth for interpreting OCT contrasts, and validation for future automated segmentation of the OCT volumes. We collected a total about 800 50-*µ*m thick slices during the four OCT imaging sessions, preserving their sequential order, and divided them into four interlaced series of about 200 slices, spaced 200 *µ*m apart. The first series was stained with thionin for Nissl body to reveal healthy neuron cell bodies and cytoarchitecture. The second used Myelin Basic Protein (MBP) immunohistochemistry to highlight myelinated fiber sheaths. The third used Tryptophan Hydroxylase 2 (TPH2) to label serotonergic neurons — a population of particular relevance to SIDS. The fourth series is reserved for protocol optimization and future use. All slices were digitized using a Keyence microscope (Keyence Corporation of America, Itasca, IL, USA; 10× objective, 1.88 *µ*m pixel size) and examined under conventional light microscope by a trained neuropathologist against published infant standards of normality (Kinney and Armstrong, 1996); no macroscopic or obvious histological abnormalities were observed.

#### 2.5.1 Neuron staining

Sections were mounted on gelatin-dipped glass slides, dried overnight, defatted in chloroform-alcohol, over-stained for three minutes in 0.05% thionin (pH 4.8), differentiated and dehydrated in ascending series of alcohol, and coverslipped from xylene with Permount. An example of a Nissl-stained sample can be found in Figure 6D.

#### 2.5.2 Myelin and serotonergic neuron staining

Free-floating fixed sections were processed in 12-well plates. Endogenous peroxidase activity was blocked using Dako Dual Endogenous Enzyme Block (Cat # S200389; Agilent, Santa Clara, CA) for 10 minutes at room temperature, followed by H_2_O washes (3 × 5 minutes). Nonspecific binding was blocked for 1 hour at room temperature with shaking in PBST (Phosphate buffered saline + 0.1% Triton X-100) with 4% horse serum. Sections were incubated overnight at 4°C with shaking in either TPH2 [1:1500] (Cat #ab121013; Abcam, Waltham, MA) or MBP [1:500] (Cat # 836504 [SMI94]; Biolegend, San Diego, CA), both diluted in PBST containing 4% horse serum. After primary antibody incubation, sections were washed (5 × 5 minutes, PBST) and incubated for 30 minutes, shaking at room temperature, with the appropriate biotinylated secondary antibody — horse anti-goat (Cat # BA-9500; Vector Laboratories, Newark, CA) for TPH2 and horse anti-mouse (Cat # BA-2000; Vector Laboratories, Newark, CA) for MBP — at 1:200 dilution. Following another round of PBST washes (5x5 minutes) at room temperature, detection was performed using the Vectastain Elite ABC-HRP kit (Cat # PK-7100; Vector Laboratories, Newark, CA). Sections were washed in H_2_O, stored in phosphate buffer at 4°C, mounted onto gelatin dipped (“subbed”) glass slides, dried overnight and then dehydrated in ascending series of alcohol and coverslipped from xylene with Permount. Examples of TPH2 and MBP stains can be found in Figure 6E and 6F, respectively.

### 2.6 Multimodal Image Registration

All modalities were registered into a common spatial coordinate space defined by the whole-brain MRI. The brainstem MRI was aligned to the whole-brain MRI and the OCT scattering coefficient volumes were aligned to the brainstem MRI. In both cases, an initial manual rigid alignment in FreeView (FreeSurfer’s visualization tool (Fischl, 2012)) provided a starting estimate, which was then refined using a robust automated registration method (Reuter et al., 2010). Normalized Mutual Information (NMI) was used as the objective function for both tasks — including the unimodal whole-brain to brainstem MRI reg- istration — because the MRI contrasts of the two datasets differ substantially across flip angles. The registration between the whole brain and the brainstem MRI was computed using a rigid transformation. Because the brainstem and cerebellum were repositioned relative to the cerebrum during the whole-brain acquisition (tucked beneath the cerebrum to achieve a tight pack), the cerebellum was masked out of the brainstem MRI before computing this alignment. The OCT scattering coefficient volumes were aligned to the brainstem MRI using an affine transformation, which accommodates voxel size inaccuracies. Registration between the OCT volumes and their corresponding histology slices requires handling the nonlinear deformations introduced by the histology process and will be addressed in future work.

## 3 Results

### 3.1 OCT Contrast

#### 3.1.1 Projection volumes

Figure 3 shows the AIP, MIP, retardance, and optic axis volumes of the caudal medulla (block 1, Figure 2). Three anatomical views are represented: axial or the plane of imaging (ventral-dorsal and left-right), sagittal (ventral-dorsal and inferior-superior), and coronal (inferior-superior and left-right). All projection volumes (AIP, MIP and retardance) can be visualized using Neuroglancer: caudal medulla, rostral medulla / caudal medulla, rostral pons and midbrain.

The four contrasts are complementary and together provide a more complete picture of brainstem organization than any single contrast alone.

In the AIP, white matter tracts appear darker than nuclei due to their higher scattering coefficient. Through-plane fiber tracts (going inferior-superior) — including the pyramids (py), cuneate tracts (cu), and spinal trigeminal tracts (sp5) — appear nearly black, reflecting near-complete absorption of backscattered light when fibers run parallel to the incident beam. The AIP and retardance are strongly anti-correlated: regions that are dark in the AIP are bright in the retardance, which maps myelin density. This anti-correlation directly demonstrates that PSOCT reliably distinguishes white matter from gray matter through optical properties alone, without reliance on myelin maturity.

The MIP reveals finer structural details not visible in the AIP, including small fiber bundles within the reticular formation. The optic axis encodes in-plane fiber orientation by color; because most brainstem tracts run through-plane, the volume predominantly reflects the optical system birefringence (green-blue hue), but in-plane tracts — including the decussation in the midline of the medial lemniscus (ml) and the fibers of the amiculum (ami) — are clearly resolved (last column of Fig. 3).

Taken together, these four contrasts demonstrate that PSOCT can delineate the major nuclei and tracts of the infant brainstem at 10 *µ*m resolution despite the low and variable myelination of the tissue — a capability that, as shown in Section 6, is not achievable with MRI at this developmental stage.

#### 3.1.2 Scattering coefficient volumes

Figure 4 shows the 20 *µ*m isotropic scattering coefficient volume of the rostral medulla and caudal pons (block 2 of Figure 2). All three anatomical views are represented: axial (v-d and l-r), coronal (i-s and l-r) and sagittal (i-s and v-d). We share the data on the DANDI platform and visualize them using Neuroglancer: caudal medulla, rostral medulla / caudal pons, rostral pons and midbrain.

**Figure 4:**
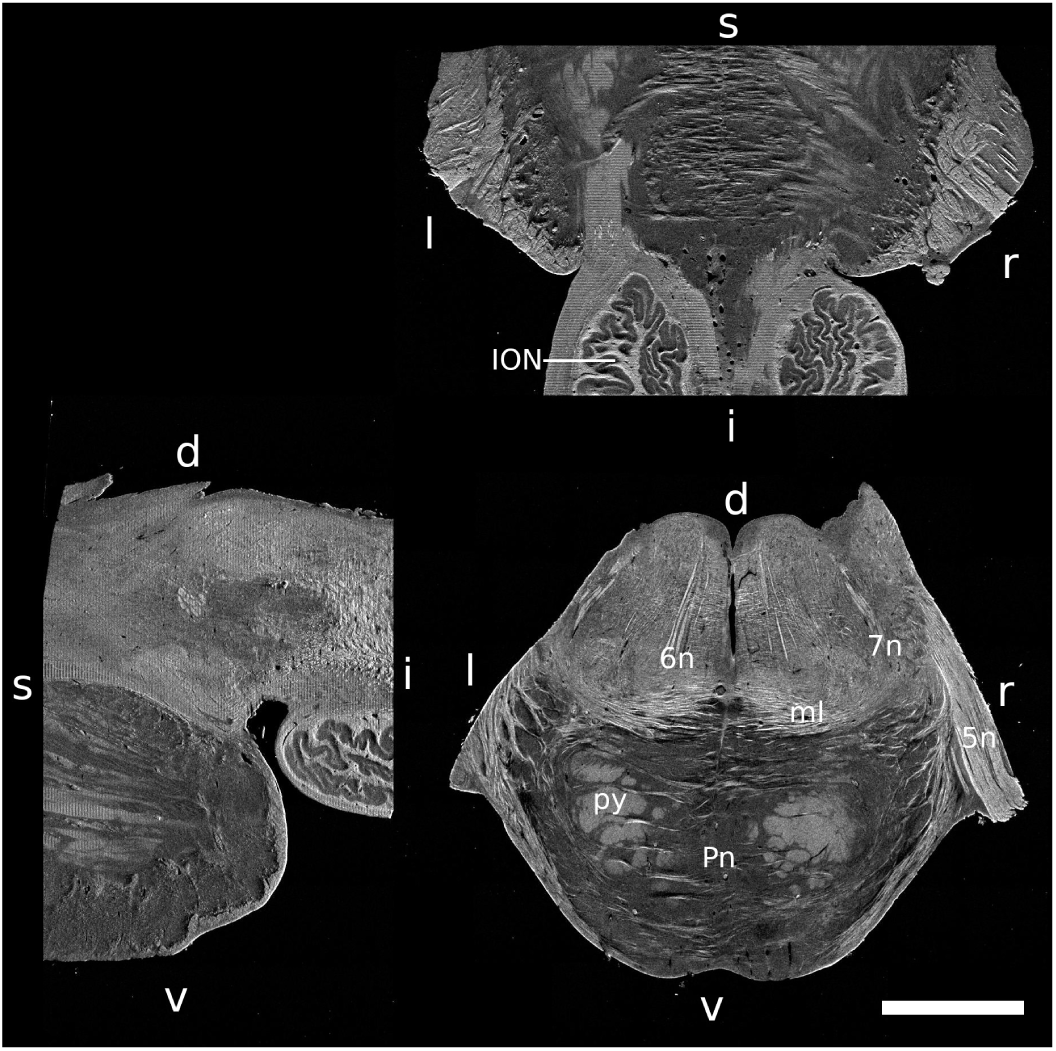
OCT-derived 20 *µ*m isotropic scattering coefficient volume of the rostral medulla and caudal pons areas (block 2 from Figure 2). **d:** dorsal, **v:** ventral, **i:** inferior, **s:** superior, **l:** left and **r:** right. 5n: trigeminal nerve, 6n: abducens nerve, 7n: facial nerve, ION: inferior olivary nucleus, Pn: pontine nucleus, py: pyramid, sp5: spinal trigeminal tract. Scale bar: 5 mm.

The scattering coefficient *µ*_s_ clearly differentiates tissue composition across the brainstem. Gray matter nuclei — including the inferior olivary nucleus (ION) and pontine nucleus (Pn) in Figure 4 — present low *µ*_s_, consistent with their cell-body-dense composition. White matter tracts present higher *µ*_s_, with myelinated fibers showing the highest values such as the pyramid (pyr) and medial lemniscus (ml). Notably, the cranial nerves — abducens (6n), facial (7n), and trigeminal (5n) — are sharply delineated by their elevated *µ*_s_, demonstrating that the scattering coefficient volume resolves individual cranial nerves within the intact brainstem block. This level of structural specificity at 20 *µ*_s_ resolution in three dimensions is not achievable with either MRI modality used in this pipeline.

### 3.2 Modality Registration

Figure 5 shows the composite registration result of our imaging pipeline across all three orthogonal planes. The whole-brain MRI, brainstem MRI, and OCT scattering coefficient volumes are displayed in a common coordinate space, demonstrating successful multi-modal alignment across a resolution range of 550 *µ*m to 20 *µ*m. The registration of both MRI datasets and all four OCT scattering coefficient isotropic volumes can be explored using this Neuroglancer scene.

**Figure 5:**
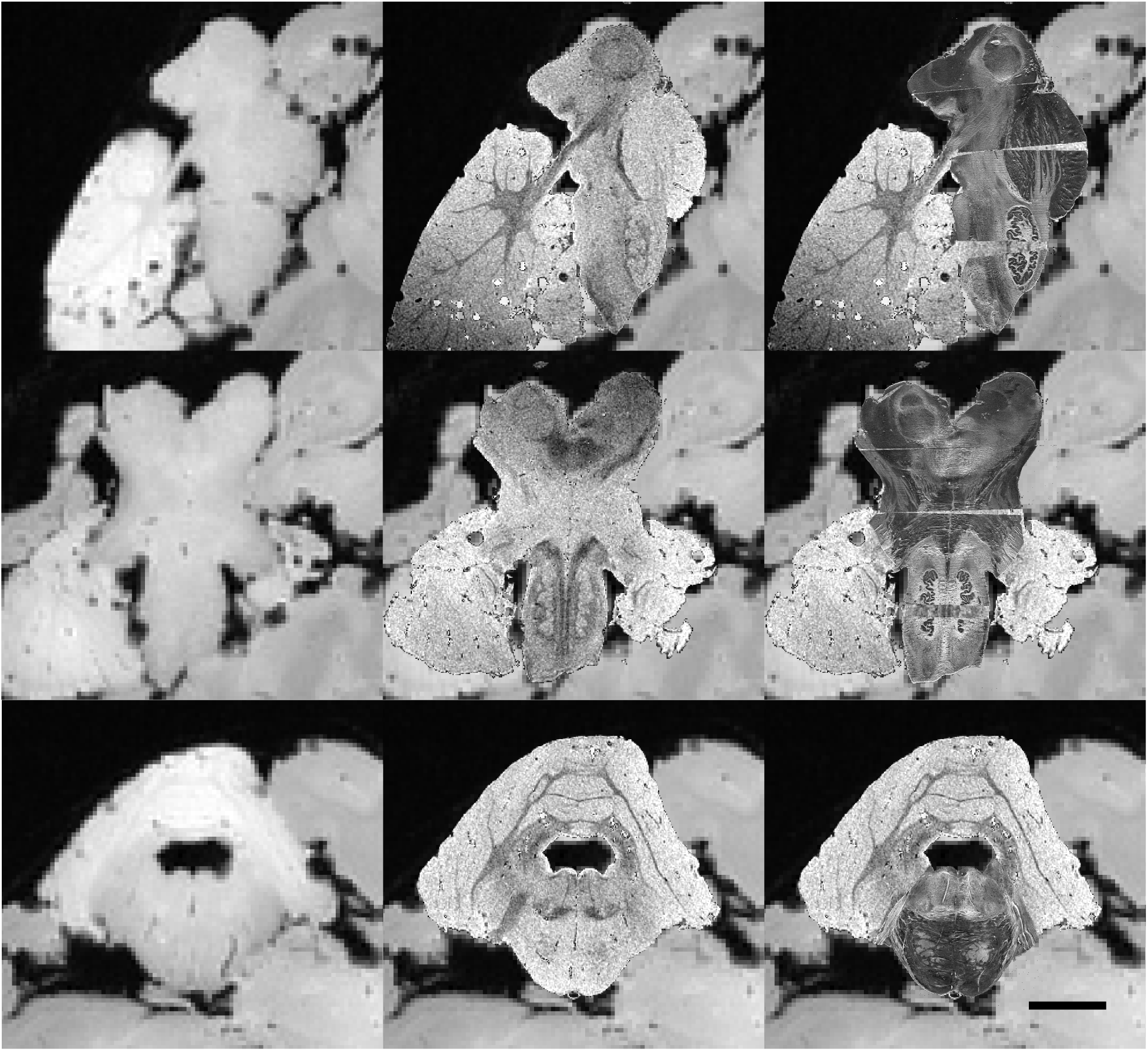
Composite image of the registration pipeline. **First column**: the whole brain MRI (550 *µ*m isotropic, multi-echo, FLASH, flip angle 30°); **second column**: the registered brainstem MRI (150 *µ*m isotropic, multi-echo, FLASH, flip angle 5°) overlaid onto the whole brain MRI; **third column**: four blocks of OCT volumes (20 *µ*m isotropic scattering coefficient) are warped into the whole brain MRI space (and displayed over both whole brain and brainstem MRI images). Three orthogonal imaging planes are displayed: sagittal (first row), coronal (second row) and axial (third row). Scale bar: 1cm.

**Figure 6:**
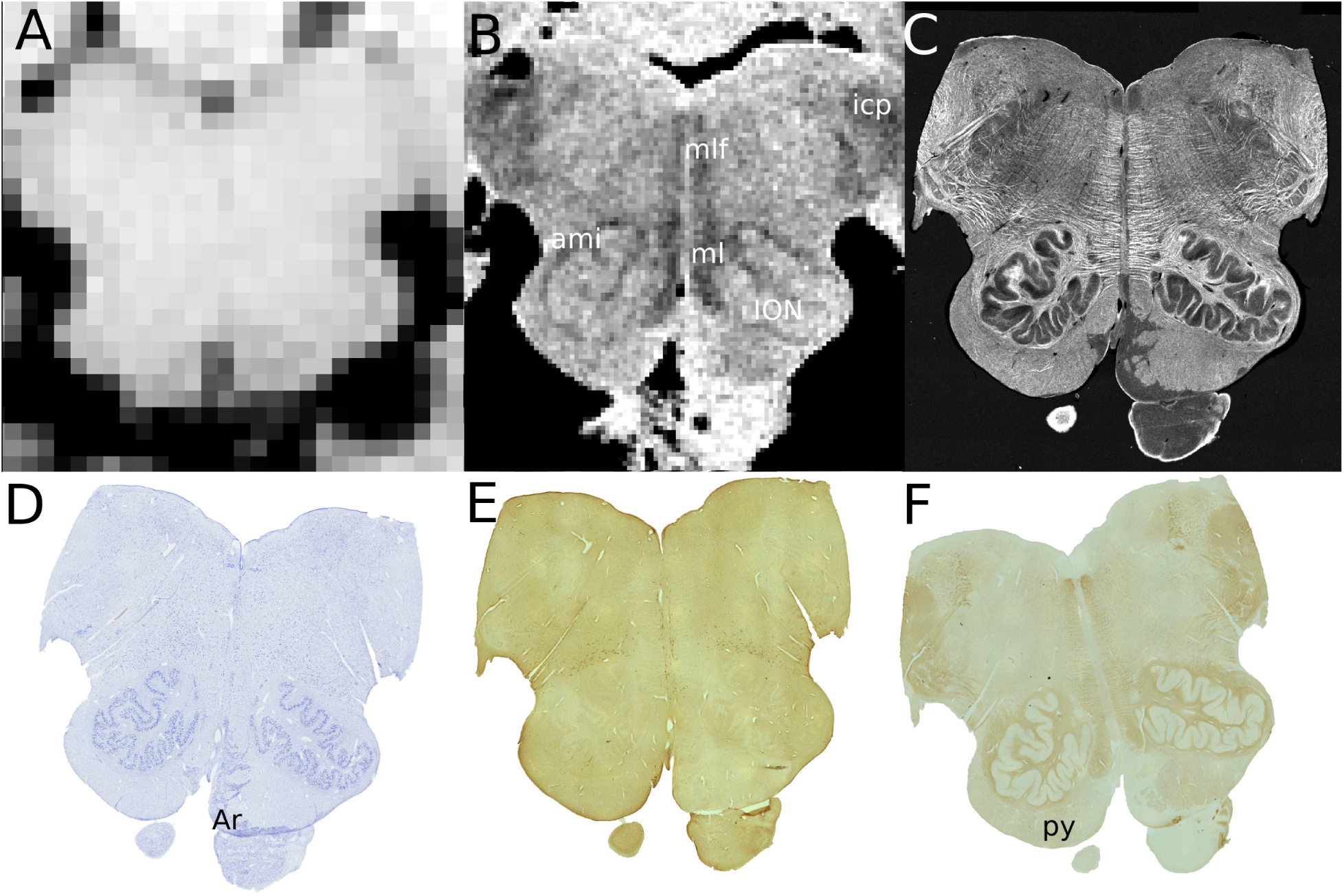
All modalities acquired at the level of the medulla: A) whole brain MRI (FLASH, flip angle 30°, 550 *µ*m), B) brainstem MRI (FLASH, flip angle 5°, 150 *µ*m), C) OCT-derived scattering coefficient (10 *µ*m), D) Nissl staining (healthy neurons), E) TPH2 staining (serotonergic neurons) and F) MBP (myelinated fibers). ami: amiculum, Ar: arcuate nucleus, icp: inferior cerebellar peduncle, ION: inferior olivary nucleus, ml: medial lemniscus, mlf: medial longitudinal fasciculus, py: pyramid.

Rigid registration between the whole-brain and brainstem MRI datasets was successful despite the repositioning of the brainstem and cerebellum relative to the cerebrum during acquisition (Reuter et al., 2010). Affine registration between the brainstem MRI and OCT volumes yielded scaling and shearing factors close to theoretical expectations across all four tissue blocks. The one exception was the midbrain block, where the scaling factor in the ventral-dorsal direction deviated from the theoretical value of 7.5 by 8.8% and shearing coefficients reached approximately 5% — modestly above the less-than-2% observed elsewhere. These deviations are consistent with a small degree of tissue deformation in the midbrain during MRI packing and do not affect the other three blocks (caudal medulla, rostral medulla/caudal pons, rostral pons). Overall, across the full brainstem volume, the registration framework successfully unifies all modalities in a shared three-dimensional space suitable for multiscale visualization and analysis.

## 4 Discussion

Human myelination in the brainstem begins in the medial longitudinal fasciculus of the medulla, the site of interconnections between the extraocular muscles, at midgestation of a term pregnancy, i.e., 20 gestational weeks of fetal life. The trajectories of myelination in different brain tracts continue and vary through fetal life, infancy, and beyond (Brody et al., 1987; Kinney et al., 1994), with molecularly mature myelination occurring only in the third decade of adulthood. Many brainstem pathways begin to myelinate in fetal life, and reach full myelination by the end of the first postnatal year; these are considered early myelinators. Other pathways begin to myelinate after birth and reach full myelination at variable time points across the second postnatal year of life as late myelinators. The very prolonged myelinators begin to myelinate before birth and continue to myelinate after the second year, e.g., the central tegmental tract carrying fibers of the brainstem reticular formation. This potentially prolonged process and the impact of lipid-bound myelin on water diffusion have significant implications for technical considerations in developmental brainstem studies with high resolution MRI.

### 4.1 PSOCT Overcomes the Principal Limitation of Infant Brain Imaging

Figure 6 presents all four modalities at the same level of the medulla. The progression from MRI to OCT to histology illustrates both the resolution gains and the qualitative contrast differences that motivate the pipeline.

The whole-brain MRI at 550 *µ*m (Figure 6A) shows very low contrast throughout the medulla. This is a consequence of two compounding factors: the limited resolution relative to the size of brainstem nuclei, and the low myelin content of the one-month-old tissue, which diminishes the primary source of white-gray matter contrast in MRI. The brainstem MRI at 150 *µ*m (Figure 6B) improves resolution 3.5-fold and reveals additional structure — including the inferior olivary nucleus (ION) — but displays the inverted contrast characteristic of unmyelinated infant tissue, where white matter tracts such as the medial lemniscus (ml), medial longitudinal fasciculus (mlf), inferior cerebellar peduncle (icp), and amiculum (ami) appear darker than the surrounding gray matter. The definition of fine structures remains limited: the convoluted laminar structure of the ION, for example, is hinted but not resolved.

The OCT scattering coefficient at 10 *µ*m lateral resolution (averaged over 50 *µ*m to match the thickness of the histology slice) (Figure 6C) resolves the same structures with markedly greater specificity. The highly convoluted shape of the ION — confirmed by the Nissl stain in Figure 6D — is clearly delineated, and additional structures not visible in either MRI dataset are revealed: the arcuate nucleus (Ar) and in-plane fiber bundles within the ml, mlf and reticular formation. This represents a genuine increase in anatomical information content, not simply a resolution improvement. The OCT achieves this without any dependence on myelination state: contrast derives from the intrinsic scattering and birefringent properties of the tissue, which are present regardless of myelin maturity. This is the central advantage of PSOCT for infant brain imaging.

The histological stains (Figure 6D–F) validate and extend the OCT findings at cellular resolution. The Nissl stain confirms the ION morphology and the presence of the arcuate nucleus seen in the OCT. The MBP stain confirms the low overall myelination of the tissue, with the highest myelin density in the icp, ami, ml, and mlf, and notably less in the pyramidal tracts (py) - a finding consistent with the known myelination timeline of these pathways (Brody et al., 1987; Kinney et al., 1988). The TPH2 stain maps the distribution of serotonergic neurons, a population of direct relevance to SIDS research. Importantly, while histology provides information that OCT cannot — cellular identity, molecular markers, and laminar cytoarchitecture — it does so only in two dimensions and with deformation introduced by the sectioning and staining process. The OCT volume provides the three-dimensional structural context within which the histological findings can be interpreted and spatially located.

### 4.2 Infant and Adult Imaging: What the Comparison Reveals

Figure 7 directly compares the infant brainstem from this study with an adult medulla (76-year-old male control) at the same MRI resolution (150 *µ*m) and the same OCT contrast (retardance), displayed on an identical intensity scale, both with in-plane resolution of 10 *µ*m. For the MRI, no postprocessing was applied. The adult scan was acquired with a field strength of 7 T and the best contrast is achieved with a flip angle of 20°. The infant scan was acquired with a field strength of 3 T and the best contrast is achieved with a flip angle of 5°. The figure reveals a pattern that has important implications for how the pipeline should be interpreted and applied.

**Figure 7:**
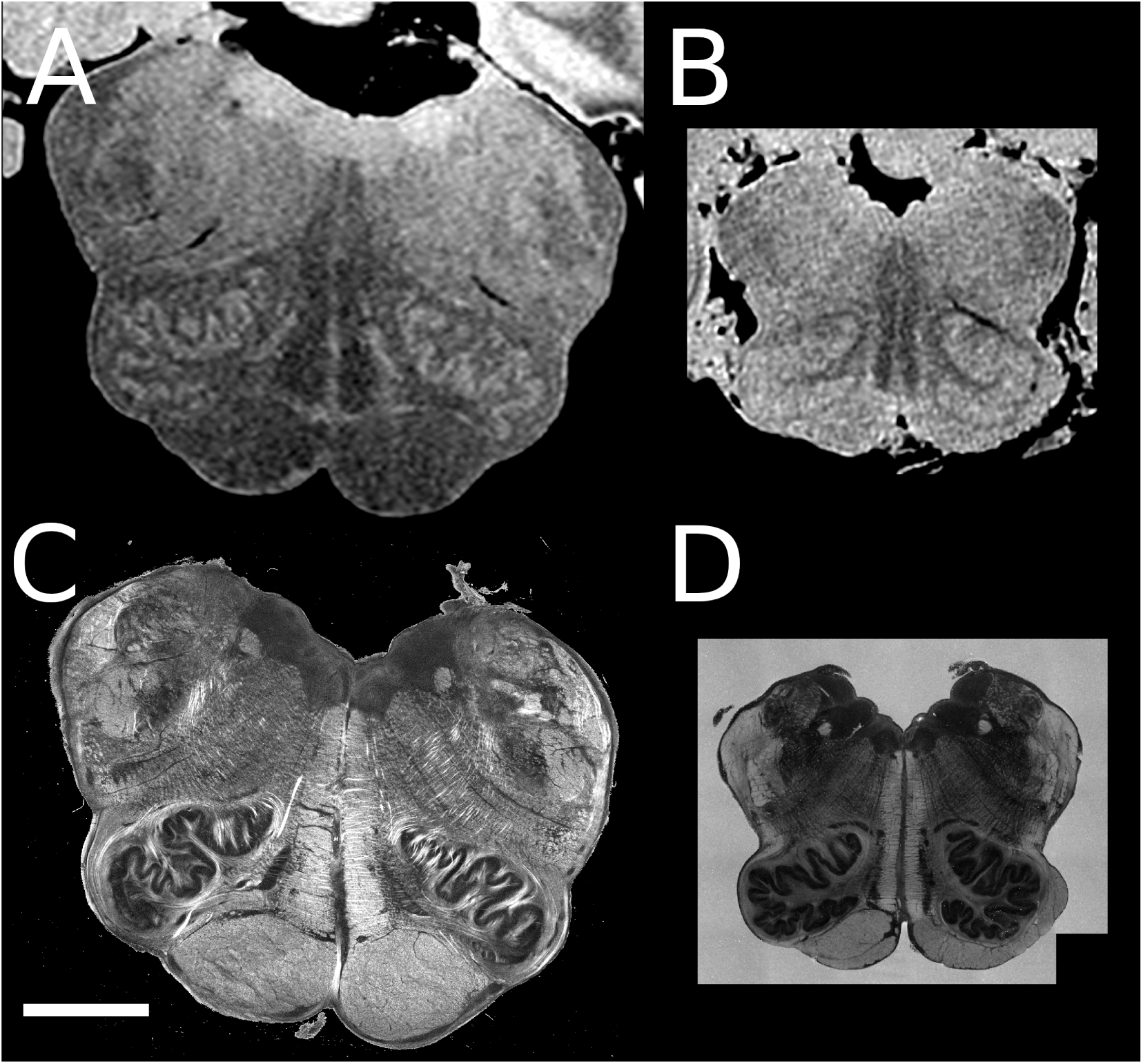
(A, C) Adult and (B, D) infant medulla imaged by MRI (both 150 *µ*m isotropic, (A) FLASH, flip angle 20° and (B) FLASH, flip angle 5°) and by OCT showing the retardance (C and D, 10 *µ*m in-plane, both on the same intensity scale). Scale bar = 5 mm.

In MRI (Figures 7A and 7B), the adult medulla shows clear contrast between white and gray matter structures, with the convoluted morphology of the ION well resolved. The infant medulla, imaged at the same resolution, shows substantially flatter contrast throughout — a direct consequence of incomplete myelination rather than any limitation of the imaging system or acquisition protocol. This is not an issue that can be resolved by improving MRI hardware or extending scan time; it reflects the fundamental tissue biology of the developing brain.

In OCT (Figures 7C and 7D), the images are qualitatively similar. Both the adult and infant datasets reveal comparable levels of structural detail, and the ION and its convoluted morphology are clearly resolved in both. The primary difference is one of signal intensity rather than structural specificity: overall retardance is lower in the infant, reflecting lesser myelin content, and the high-retardance fiber bundles visible as bright striations in the adult — around the ION and within the reticular formation — are absent or minimal in the infant. However, this reduction in signal intensity does not prevent structural delineation. The boundaries between nuclei and tracts remain visible in the infant OCT data because PSOCT contrast is not binary with respect to myelination — it is sensitive to any degree of birefringence in the tissue, including the partial myelination present at one month of age.

This comparison highlights the fact that the challenge of imaging the infant brainstem is not simply one of scale but of contrast mechanism. PSOCT addresses the contrast problem directly, and the adult-infant OCT comparison demonstrates that the pipeline delivers structurally informative data across the myelination spectrum. This has direct implications for applying the pipeline to developmental studies spanning fetal life through early childhood, and to pathological cases where myelination may be further disrupted.

### 4.3 Scope and Limitations

Several limitations of the current pipeline should be noted. First, the specimen presented here is a single normative case; while this is appropriate for a proof-of-concept demonstration, validation across a larger sample — including cases with confirmed brainstem pathology — will be necessary to establish the pipeline’s sensitivity to cellular and molecular abnormalities. Second, registration between OCT volumes and their corresponding histology slices remains an open problem. Histological processing introduces nonlinear deformations — tissue shrinkage, tearing, and folding — that rigid and affine registration cannot fully correct. Developing a robust volumetric histology reconstruction method using the OCT volume as a deformation template is a priority for future work. Third, non-linear registration between the both MRI dataset and OCT to take into account some local deformations such as the ones introduced by packing for MRI imaging and embedding for OCT imaging will be explored to improve further the alignement between all modalities. Fourth, diffusion MRI data were acquired for both the whole brain and brainstem but are not reported here; incorporating these data into the registered framework will add white matter tractography as a further modality bridging the scales between MRI and OCT.

## 5 Conclusion

Brainstem disorders such as SIDS have resisted neuroimaging-based investigation for decades because the abnormalities they produce — cellular, serotonergic, molecular — occur at scales and with contrast properties that conventional MRI cannot access, particularly in the immature infant brain. The pipeline introduced here directly addresses this gap.

By integrating whole-brain MRI, brainstem-specific MRI, PSOCT, and histology with immunohistochemistry in a common three-dimensional coordinate space, the pipeline spans four orders of resolution mag- nitude — from 550 *µ*m to 1.88 *µ*m — while preserving spatial correspondence across all modalities. The central demonstration of this paper is that PSOCT provides reliable, high-resolution tissue contrast in the infant brainstem independent of myelination state, resolving nuclei and tracts at 10 *µ*m in three dimensions where MRI produces ambiguous or inverted contrast at 150 *µ*m. Histology validates and extends these findings at cellular resolution, and the adult-infant OCT comparison establishes that the pipeline delivers structurally informative data across the full spectrum of myelination states encountered in developmental neuroanatomy. Our imaging pipeline will also allow to further our understanding in the sequence of myelination of the brainstem throughout the early development.

Applied to a normative 34-day-old infant brainstem as a proof of concept, the pipeline is now positioned for application to cases where brainstem pathology is the primary question. The framework is generalizable to any region of the developing brain — including the cerebellum, hippocampus, thalamus, and cortex — and to other developmental stages. Its greatest potential impact, however, may be in the context that motivated its development: brainstem disorders such as SIDS, where the target of investigation is cellular and molecular rather than gross anatomical, and where the combination of three-dimensional structural context from PSOCT with molecular specificity from immunohistochemistry offers capabilities that neither modality provides alone.

## 6 Data Sharing

We publicly share all imaging data presented at: https://dandiarchive.org/dandiset/000874/draft

## Funding

This work was supported by the National Institute of Health (NICHD 5R01HD102616, 5R21HD095338, 1R21NS106695-01A1), the American SIDS Foundation and Chan-Zuckerberg Initiative DAF, an advised fund of Silicon Valley Community Foundation (2019-198101 and 2021-244261). This research was also made possible by the resources provided by National Institutes of Health P41RR014075 and shared instrumentation grants 1S10RR023401, 1S10RR019307, and 1S10RR023043.

## Declaration of Competing Interests

The authors have nothing to declare.

## Acknowledgments

First and foremost, we would like to thank and acknowledge the family who donated brain tissue for our research. We would also like to acknowledge the hard work and many hours spent on anatomical labeling by Sam Blackman and Seoyoon Kim, two undergraduate students at the time of our investigation.

## Notes

https://dandiarchive.org/dandiset/000874/draft

### Competing Interest Statement

The authors have declared no competing interest.

